# Volumetric Printing across Melt Electrowritten Scaffolds Fabricates Multi-Material Living Constructs with Tunable Architecture and Mechanics

**DOI:** 10.1101/2023.01.24.525418

**Authors:** Gabriel Größbacher, Michael Bartolf-Kopp, Csaba Gergely, Paulina Nuñez Bernal, Sammy Florczak, Mylène de Ruijter, Jürgen Groll, Jos Malda, Tomasz Jüngst, Riccardo Levato

## Abstract

Major challenges in biofabrication revolve around capturing the complex, hierarchical composition of native tissues. However, individual 3D printing techniques have limited capacity to produce composite biomaterials with multi-scale resolution. Volumetric bioprinting recently emerged as a paradigm-shift in biofabrication. This ultra-fast, light-based technique sculpts cell-laden hydrogel bioresins into three-dimensional structures in a layerless fashion, providing unparalleled design freedom over conventional bioprinting. However, it yields prints with low mechanical stability, since soft, cell-friendly hydrogels are used. Herein, for the first time, the possibility to converge volumetric bioprinting with melt electrowriting, which excels at patterning microfibers, is shown for the fabrication of tubular hydrogel-based composites with enhanced mechanical behavior. Despite including non-transparent melt electrowritten scaffolds into the volumetric printing process, high-resolution bioprinted structures were successfully achieved. Tensile, burst and bending mechanical properties of printed tubes were tuned altering the electrowritten mesh design, resulting in complex, multi-material tubular constructs with customizable, anisotropic geometries that better mimic intricate biological tubular structures. As a proof-of-concept, engineered vessel-like structures were obtained by building tri-layered cell-laden vessels, and features (valves, branches, fenestrations) that could be resolved only by synergizing these printing methods. This multi-technology convergence offers a new toolbox for manufacturing hierarchical and mechanically tunable multi-material living structures.

## INTRODUCTION

In the quest to restore the function of damaged tissues, additive manufacturing technologies are continuously opening new avenues to better capture the complex composition and function of native biological architectures.^[1]^ A central characteristic of biofabrication techniques is the ability to perform the automated and accurate simultaneous placement of living cells and materials (together also termed bioinks)^[2]^ in custom-designed patterns. Fabrication can be performed by means of light-based printing, such as stereolithography and digital light processing,^[3, 4]^ extrusion-based,^[5, 6]^ droplet-based or inkjet printing,^[7–9]^ and fiber reinforcing technologies.^[10, 11]^ All these technologies have their individual benefits and challenges with respect to resolution, shape-fidelity, cell-viability, compatible ranges of materials (inks or resins), and printing/processing time. Yet, while each technique excels in processing certain subsets of inks and object geometries, native biological tissues are characterized by their unique multicellular and multi-material composition, shape and hierarchical architecture, with features from the submicron to the macro-scale. Most importantly, both mechanical and biological function are intimately correlated to this multi-scale, multi-material hierarchy.^[1]^ New directions in the biofabrication field hold the promise to bridge this gap by converging different (and previously incompatible) bioprocessing technologies,^[12, 13]^ with the aim to produce engineered tissue constructs that, by combining their benefits and range of achievable prints, better mimic salient features of their native counterparts, to eventually restore and replace damaged tissues.

Volumetric bioprinting (VBP) is a recently developed biofabrication technology to sculpt hydrogels into free-form 3D structures.^[14]^ In VBP, a hydrogel with the addition of a photocrosslinking agent is placed in a rotating platform and a light source (i.e., a laser), in combination with a spatial light modulator (such as a digital micromirror device, DMD), is subsequently used to deliver a sequence of filtered tomographic back-projections onto this volume. The sum of the different projections rapidly generate an anisotropic, 3D light-dose distribution within the build volume, thereupon activating the polymerization reaction only in correspondence to the desired object.^[15]^ This process is extremely fast (<30 seconds to produce several cm^3^ parts)^[16]^ compared to conventional extrusion-based printing (20 min), while offering equal or greater resolution (40 - 200 μm) with higher spatial freedom.^[14, 15]^ Although promising, the materials used with volumetric printing are hydrogels, which are intrinsically soft, while most tissues in the body also need to account for mechanical stability and load bearing capacity. Techniques to mechanically stabilize soft hydrogels include the formation of interpenetrating polymer networks, the inclusion of nanoparticles, or the inclusion of fibrous reinforcements, such as structures produced via fused deposition modelling, electrospinning, and melt electrowriting (MEW).^[17]^ In particular, MEW generates (sub)micrometer-scale fibers by applying a high voltage to the polymer melt. Fibers are deposited onto a moving collector plate, allowing for control over fiber deposition and subsequent scaffold architecture.^[17, 18]^ Scaffolds made with MEW have proven to facilitate cell alignment,^[19]^ and increase the mechanical stability of tissue engineered constructs.^[20]^ This technique is compatible with printing a variety of thermoplastic materials,^[21, 22]^ whose biochemical composition can further be modified post-writing with surface coatings.^[11,21,23]^ Recently, first steps have also demonstrated the possibility to print hydrogel-based fibers.^[24]^ Notably, fiber deposition can be done on a flat collector plate but also on a mandrel for the fabrication of intricate tubular scaffolds.^[25–27]^ Tubular geometries are of particular interest for biological applications, as they recur in many tissues, and can therefore be applied as scaffolds for blood vessels, airways, intestinal and tubular kidney structures,^[28]^ among others.^[26, 27]^

One of the drawbacks of MEW tubular scaffolds is that the inclusion of cells is generally done post-printing, either by direct seeding on top of the fiber strands, or by casting with a hydrogel carrier.^[29–31]^ Promising composite structures have been produced, exploiting the unique ability of MEW to provide mechanical reinforcement to other cell-carrying materials.^[32]^ However, since the cell-laden compartment can only be loaded in simple geometries following the electrowritten mesh pattern, replicating complex, branched and tortuous geometries typical of native tissues remains challenging.

In this work, we demonstrate for the first time the convergence of volumetric bioprinting with melt electrowriting, to pattern multiple materials and cell types in any custom-desired geometry even within opaque polymeric microfibrous thermoplastic meshes. In shaping these novel hydrogel-cell-microfiber composites, as a proof-of-concept to demonstrate the applicability and versatility of this technology, we built a broad array of constructs that mimic key features of native blood vessels. The intricacy in design ranges from double-branched structures to multi-cellular and fenestrated, structurally reinforced scaffolds with tunable mechanical properties, and cell-laden architectures not possible with previously existing techniques alone.

## RESULTS AND DISCUSSION

The convergence of different 3D printing technologies has become a widely studied concept in the fields of biofabrication and tissue engineering, given the potential to exploit the advantages of different technologies to combine different classes of materials in a single object, and to create living, hierarchical structures.^[12]^ In the present study, we aim to elucidate the potential and advantages offered by the convergence of volumetric (bio)printing (VP), which allows the fabrication of highly complex, centimeter-scale structures using hydrogel-based bioresins,^[14]^ and melt electrowriting (MEW), which allows the creation of highly organized fiber architectures having micron-scale resolution, uniquely able to confer outstanding mechanical properties to low-stiffness hydrogels. To accomplish this, tubular MEW scaffolds were first fabricated from poly-ε-caprolactone (PCL) and incorporated into a volumetric printing vial, which in turn was exposed to the tomographic light projections required to form a 3D object in tens of seconds (Figure 1A). As platforms to investigate the converged technique, termed VolMEW for simplicity of reference, materials widely used in the field of biofabrication for both technologies were employed in this study: PCL for MEW, given its previously established superior printing properties and medical grade nature,^[21]^ and gelatin methacryloyl (gelMA) as a bioresin for VP, which has been shown to be a fast, high-resolution choice for this printing approach.^[14, 33]^ In the present study, different MEW mesh architectures were successfully fabricated, with custom-designed i) pore shapes (i.e., rhombic with 34° and 70° winding angles and hexagonal (Figure 1Ai, B)) and ii) thicknesses (i.e., 20, 30, 40, and 60 printed layers). These tubular scaffolds were first placed into volumetric printer vials and guided in through a carbon fiber rod with controlled alignment (Supplementary Figure S1). Subsequently, the vials were loaded with gelMA. Due to the thermal gelation behavior characteristic of gelMA, the MEW mesh could be secured into its intended location and alignment even after removal of the support rod (Figure 1Aii -iv). With the MEW scaffold aligned within the printing vial, the VP process was conducted (Figure 1Av, C).

**Figure 1:**
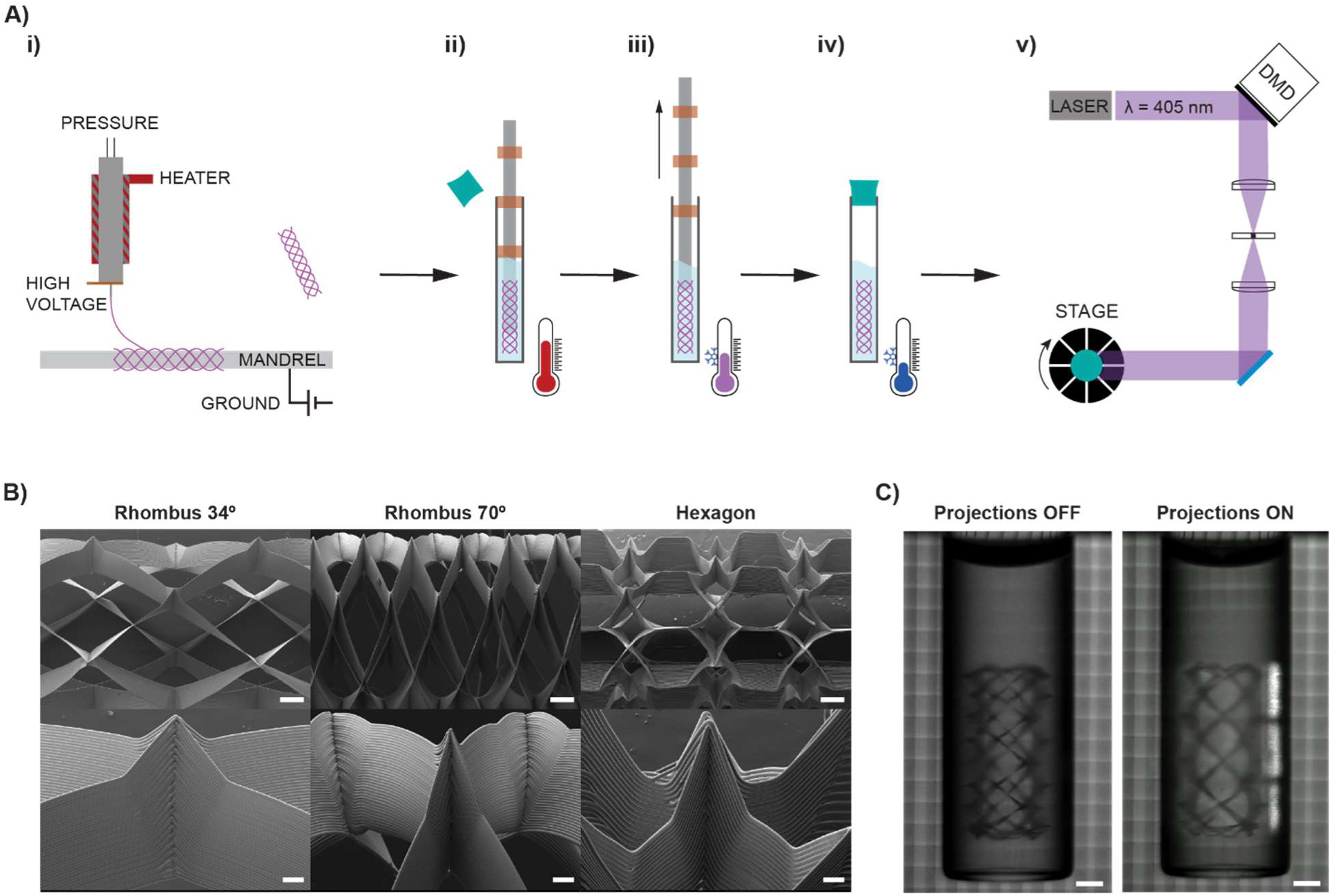
Convergence of MEW and VBP processes - VolMEW. A) Graphical overview of i) the fabrication of tubular melt electrowritten scaffolds on a rod and their subsequent incorporation into the VBP process by ii) first placing the MEW meshes in a vial of preheated gelatin methacryloyl solution using a carbon fiber rod with centering guides. iii) once the MEW tube is completely infused, the gelMA solution is gradually cooled as the mandrel is retracted without disrupting the gelling resin or the MEW mesh structure, iv) the vial is then fully gelled in ice water and v) placed in the volumetric bioprinter. B) SEM images of three different pore structures of the tested MEW tubes (scale bars = 500 µm (top panels) and 100 µm (bottom panels). C) Digital images of the VBP vials containing gelMA and a fixed, centered MEW mesh when inserted into the printer and when light projections are initiated. (scale bar = 2mm)

After establishing a simple and consistent method for setting up a hybrid VolMEW process, the effect of the opaque PCL meshes on the tomographic illumination imparted by VP was investigated to ensure the achievement of high-resolution prints (Figure 2). It was essential to first characterize this effect because the VP process relies on the undisturbed passage of tomographic light projections through the entire bioresin vat in order to induce specific photocrosslinking in the target regions. The presence of elements that attenuate or refract the light beams incoming from the spatial light modulator could therefore impair printing resolution. While the presence of scattering elements (i.e., cells, particles) can be mitigated mixing the bioresin with refractive index matching compounds,^[33]^ the effect of opaque structures had yet to be investigated. Previous research has demonstrated the possibility to perform volumetric prints around non-transparent objects, such as metallic rods. However, such prints were only achieved with bulky structures fully encasing the abovementioned rod and lack the fine features characteristic of volumetrically printed objects.^[34]^ In this study, the opacity of the PCL mesh provided a light attenuation effect by partially blocking the projection path in the regions where the PCL meshes align with the beam (occlusion points).

**Figure 2:**
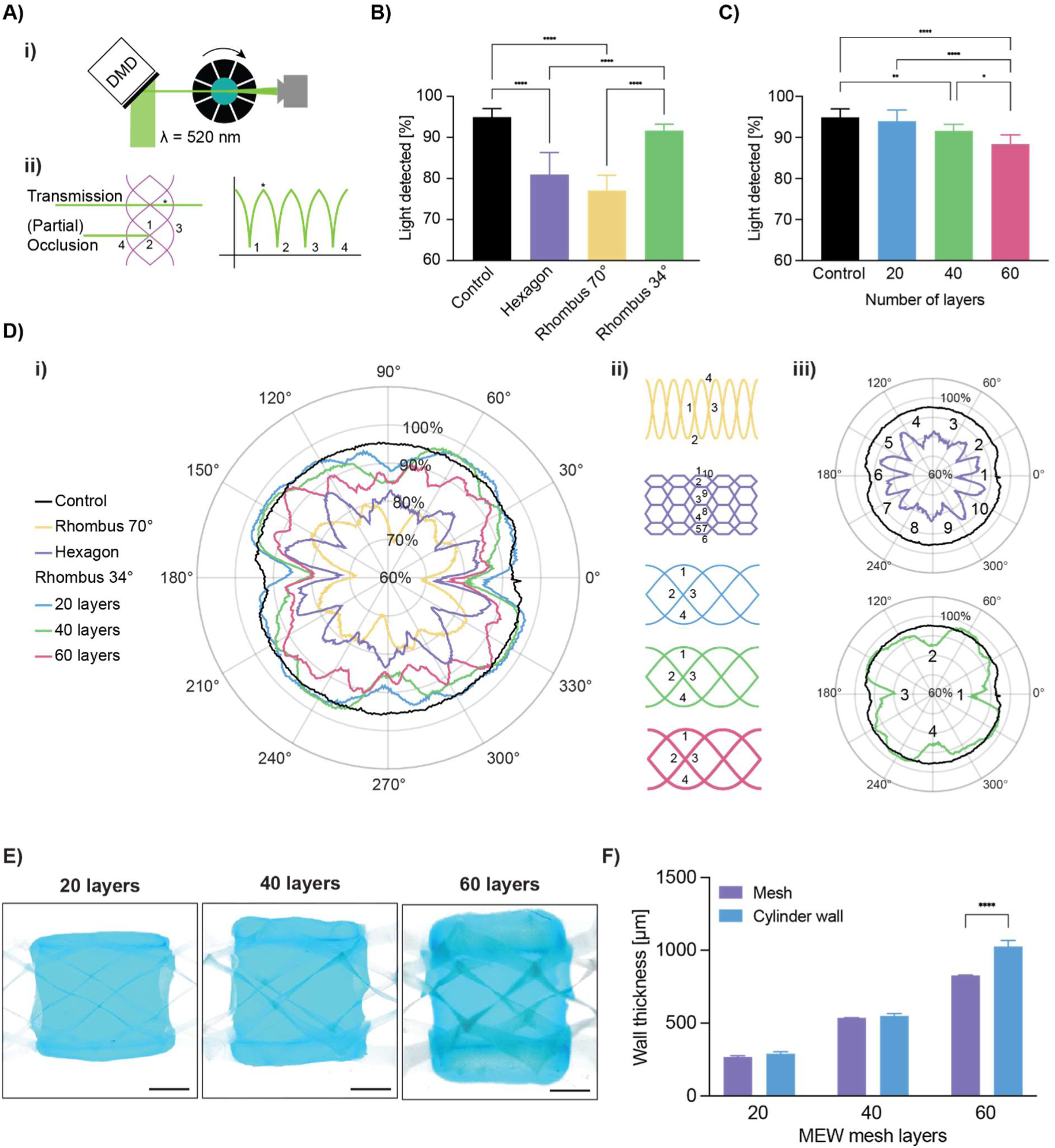
Characterization of the effect of MEW tube incorporation on beam path homogeneity through gelMA-filled VBP vials. A) Graphical overview of the test setup used to evaluate the light attenuation effect of different architectures/layers of MEW meshes. i) A 520nm laser beam is used to project a pixel array through a VBP vial containing gelatin and different MEW tube structures for one 360° rotation while a detector placed at the opposite end of the vial is captures the percentage of light passing through the vial at each angle of rotation. ii) Diagram of the numbered occlusion locations in a 34° rhombus, where the highest light attenuation is observed with an example transmission point (*) and a corresponding graph. Based on the resulting light detection measurements per structure showing the percentage of light passing through the vial, the area under the curve is plotted for B) different mesh architectures: hexagons, rhombi with 70° and 34° winding angle and mesh-free controls, as well as for C) the 34° winding angle rhombus with different number of layers (20, 40 and 60). Di) Time synchronous average percentage of light detected for the different mesh architectures and layer heights: hexagons (40 layers), rhombi with 70° (40 layers) and 34° (20, 40, 60 layers) winding angles and mesh-free controls (n=4). ii) Architectural mesh diagrams and their respective occlusion points. Each occlusion mesh point (represented by the troughs in the graph) is numbered and iii) examples are given for the corresponding graphs. E) Stereomicroscope images of VBP-printed tubes accurately encapsulating tubular MEW meshes of 20, 40 and 60 layers respectively (scale bars = 1 mm). F) Average wall thickness of the MEW meshes with different number of layers (20, 40 and 60) and the average wall thickness of the printed hydrogel surrounding these structures (n = 3, scale bars = 1 mm). * = p < 0.05, ** = p < 0.01, **** = p < 0.0001.

A custom-made setup to evaluate the light attenuation effect was devised. A laser (520 nm) was used to project a small pixel matrix through the center of a rotating VP vial containing the constructs with varying MEW pore architectures, or constructs with the same mesh geometry, but different layer heights. A photodetector was then placed on the opposite side of the vial, to determine the profile of the light beam passing through the hydrogel-soaked MEW mesh (Figure 2A, Supplementary Figure S2). The percentage of projected light intensity passing through each angle of the vial was calculated by integrating the normalized signal over a full rotation and represents the average percentage of light detected with MEW scaffold attenuations of different pore architectures (Figure 2B) and layer heights (Figure 2C). Compared to hydrogel-only control conditions, where 94.9 ± 2.0 % of light intensity could be detected after passing through the bioresin vial, all three MEW pore architectures: hexagons (80.9 ± 5.3 %) and rhombi with 70° (77.1 ± 3.7 %) and 34° (91.6 ± 1.6 %) winding angles, showed significantly lower percentages of detected light. As for the differences between architectures, the 34° rhomboid-shaped pore architecture showed the lowest attenuation effect, allowing significantly higher amounts of light passage compared to the hexagons and the 70° rhomboid. The light detection plots (Figure 2Di, Supplementary Figure S3) clearly visualize that the different architectures exhibit a series of steep dips where the light is partially blocked by each occlusion point (Figure 2Dii). Although the occlusion points coincide with a high local light attenuation, the average light passing through the vial is observed to be correlated with the pore size of the scaffold rather than the amount of total occlusion points per revolution. This hypothesis is supported by the significant difference in total percent transmitted light between 34° and 70°, where both have only four occlusion points per revolution, but the 34° rhomboid (8 pivot points) has a pore size of 4.12 mm², whereas 70° rhombi (8 pivot points) have a pore size of only 1.01 mm².^[25]^ Using the same setup, 34° rhombi were selected to investigate the effect of layer height based on the observed least significant light attenuation for this architecture. Here, compared to the mesh-free control, the 20-layer scaffold (94.0 ± 2.7 %) did not significantly decrease the amount of transmitted light, as opposed to what was found for the 40-(91.6 ± 1.6 %) and 60-(88.4 ± 2.2 %) layer structures (Figure 2C). The observed differences in light attenuation are likely caused by i) the increase in total electrowritten material, combined with ii) minor stacking errors over the entire layer height resulting in a larger occlusion zone. In addition, inherent to MEW on a rotating mandrel, each deposited layer could suffer from a minute layer shift, effectively creating a sloped and, therefore, thicker wall. Consequently, as this thicker wall rotates out of the field of view of the projected spot, it partly overcasts the detected light over a longer angular distance when compared to thinner PCL walls (Supplementary Figure S4).

Having shown that the degree of light attenuation can be controlled by pore architecture and layer height selection, the effect of shading on VP printing accuracy was assessed using 34° rhombic tubes at the different layer heights tested above. As a benchmark assay, we volumetrically printed cylinders with arbitrarily designed wall sizes to match the thickness of the MEW mesh (t_print_ = 14.8 s). All three MEW-mesh wall thicknesses (20, 40, and 60 layers) were completely encapsulated within VP-printed gelMA cylinders of equal programmed thickness as the MEW scaffolds (Figure 2E). The thickness of the wall enveloping the 20- and 40-layer scaffolds (291 ± 13 µm and 550 ± 16 µm, respectively) did not significantly differ from their respective target designs (MEW mesh thicknesses) (267 ± 10 µm and 537 ± 1 µm for the 20- and 40-layer meshes, respectively), suggesting that the potential effect of light attenuation for these scaffolds can be easily avoided with the selected volumetric printing light dose settings (Figure 2F). However, this was not observed for the 60-layer scaffold, where the resulting wall thickness encapsulating this structure (1026 ± 41 µm) was significantly larger than the mesh itself (828 ± 3 µm) (Figure 2F). The significant decrease in printing accuracy is underpinned by the light attenuation observations described previously, as this scaffold had the most noticeable effect on the passage of light through the VP sample. It is hypothesized that the thicker scaffold experiences a higher sum of diffuse backscatter reflected from the opaque PCL fibers, resulting in lower precision in delivering the light dose along the projected pattern. Consequently, off-target regions of the bioink adjacent to the MEW mesh also reached the photopolymerization threshold. Despite this effect seen in the thicker MEW samples, it is noteworthy that all three mesh thicknesses allowed VP to occur and produced homogeneous cylindrical prints. Moreover, if thicker MEW scaffolds are required for specific applications, this information on the light attenuation profile could feed future algorithms to computationally correct the tomographic projections.

In previous research focused on the VP technique for bioprinting applications, biocompatible bioresins obtained from hydrogel-based materials with relatively low mechanical stability have been used. On the one hand, these classes of hydrogels are desirable when it comes to maintaining cell viability and facilitating cell-to-cell communication.^[14,33,35,36]^ On the other hand, these bioresins are often not strong enough to withstand the harsh mechanical environment found in native tissues, be it shear, tensile, or compressive forces. The VolMEW approach has the potential to overcome this challenge and enhance the mechanical stability of the printed structures, enabling a range of tissue-specific load-bearing applications in line with the mechanical stabilization that has been achieved when MEW was combined with extrusion-based bioprinting.^[32, 37]^ To determine the mechanical properties of the hybrid VolMEW constructs, the burst and tensile performance of the composite prints were tested.

First, the burst pressure of the tubes was assessed by connecting the VolMEW tubes to a custom-made test setup via microfluidics adapters and commercially available tubing (Figure 3A). Different from what was experienced with reinforced tubes, simply mounting constructs made from non-reinforced gelMA constructs to the adapters was difficult, as these prints tended to rupture while being secured in place. Vaseline was injected into the luminal cavity of the tubes and the pressure was recorded until failure. A significant difference in burst pressure was recorded for the different geometries, which displayed increased strength when compared to the gelMA-only controls (Control: 0.66 ± 0.32 bar / Hexagon: 1.14 bar ± 0.10 bar / 34°: 1.3 ± 0.26 bar / 70°: 1.58 ± 0.17 bar)(Figure 3B). Overall, these results corroborate previous observations of hydrogel-infused tubular MEW scaffolds, which also showed enhanced tensile properties and resistance to flow-based pressures.^[31]^ From a physiological perspective, the measured values are above the maximum arterial pressure, but below expected burst strength recordings for native vascular systems.^[38]^ Nevertheless, the measurements reveal the possibility of modifying these parameters by adapting the reinforcement geometry of the hybrid construct and adjusting the properties of the hydrogel via its composition. Moreover, it should be noted that, in a tissue engineering application, cells could be added to the printed scaffold prior to implantation, and the constructs could be subjected to an in vitro maturation step to finally match the desired mechanical properties for in vivo utilization.^[39]^

**Figure 3:**
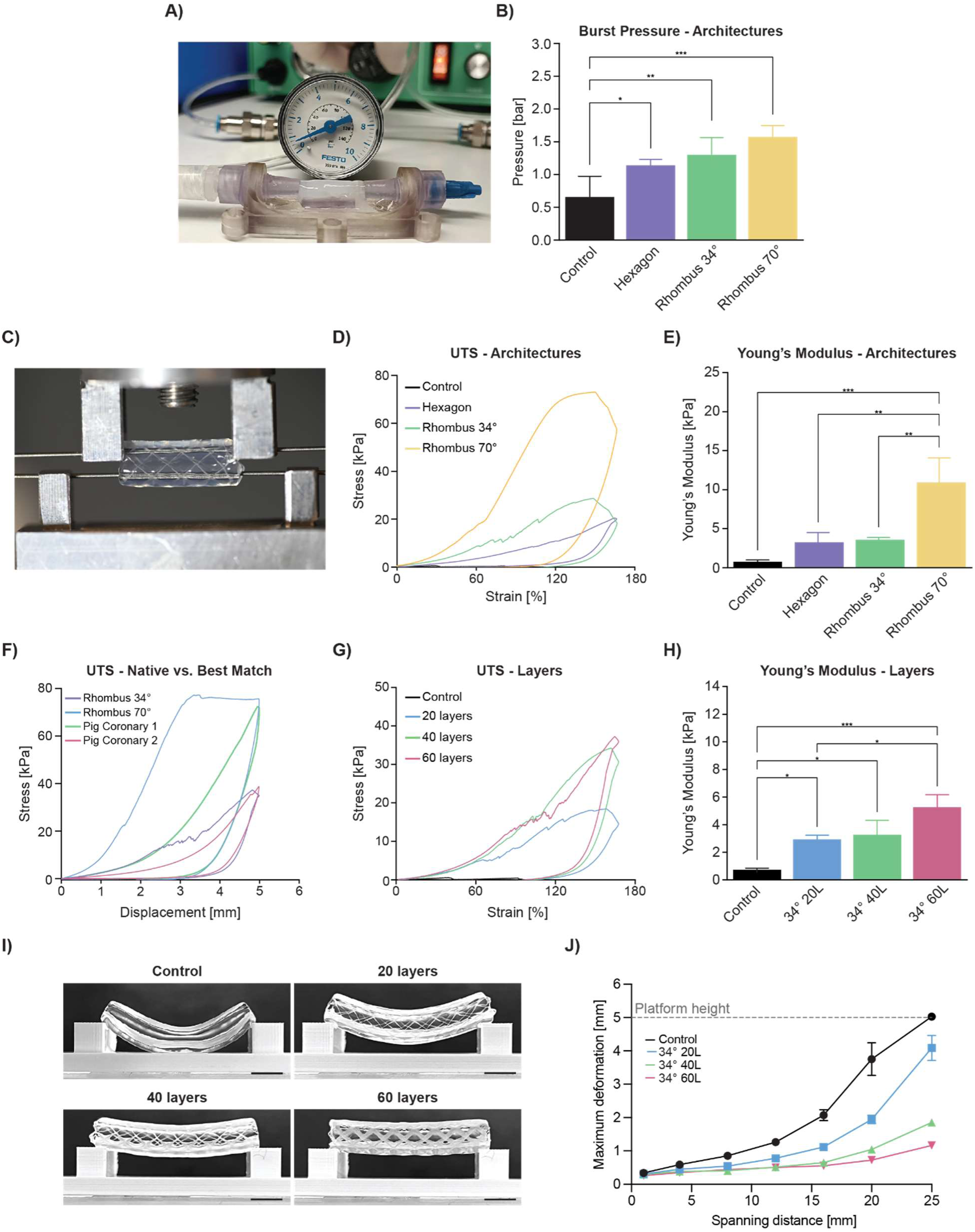
Mechanical properties of VolMEW constructs with different mesh architectures and number of layers. A) Photograph of the burst pressure test setup with the construct mounted and pressurized using part of a custom 3D printed bioreactor assembly. B) Burst pressure evaluation between different MEW reinforcement geometries (30 layers). C) Photograph of the customized mounting setup on the uniaxial tensile testing machine with the VolMEW construct in place. D) Ultimate tensile stress analysis for architectural differences between the tested MEW geometries (30 layers) within the VBP tubes. Constructs were displaced to 5 mm, corresponding to 166% strain. E) Graphical depiction of Young’s Moduli compared between MEW geometries (30 layers). F) Comparison of VolMEW construct UTS measurements with porcine coronary artery measurements obtained using the same tensile test setup and parameters. G) Ultimate tensile stress analysis of different layers of MEW construct reinforcement H) Graphical representation of Young’s Moduli compared between different layers of MEW construct (34° rhombus geometry). I) Bending resistance was assessed by placing a volumetrically printed gelMA tube reinforced with 34° winding angle rhombus MEW meshes of different number of layers (0, 20, 40 and 60) on two 5 mm high columns spanning 25 mm in length to evaluate the maximum deformation of the printed construct (scale bars = 5 mm). J) Plotted average maximum deformation of control samples without MEW reinforcement and tubes reinforced with 34° winding angle rhombus MEW meshes of different number of layers (0, 20, 40 and 60; dotted line represents the maximum height of the columns on which the constructs were placed). * = p < 0.05, ** = p < 0.01, *** = p < 0.001. n = 3 for mean values, unless indicated otherwise.

Next, a custom two-pin setup was mounted on a uniaxial tensile testing machine capable of cyclic motion and samples were mounted by inserting the pins through the lumen of the construct to perform radial tensile tests (Figure 3C). In this fashion, a motion similar to the dilation of blood vessels during systole could be replicated and recorded. This is a relevant parameter to quantify, because the compliance of a tubular construct is associated with the likelihood of graft occlusion and ultimately rejection, especially for small-diameter vascular structures.^[40]^ Another advantage of testing circular constructs in radial tensile experiments is that it more closely approximates physiological mechanical properties compared to the planar axial method of cutting the tube longitudinally and clamping it onto the testing machine.^[41]^ Conversion to stress and strain values were calculated using formulas previously used in wire myography.^[42]^ In order to evaluate the tensile performance of the composite materials, samples were strained to 5 mm, which equates to 166% elongation, a value chosen as a reference point as it exceeds physiological vascular strain levels. Figure 3D illustrates the behavior of VolMEW constructs with different reinforcement geometries compared to unreinforced constructs. Each of the reinforcement geometries showed a different hysteresis curve over cyclic loading, and all of them allowed the VolMEW constructs to withstand a full cycle of stress loading, that instead resulted in failure for VP-only (gelMA-only) constructs. A significant overall increase was shown in the recorded peak stress values for all reinforcement strategies, with hexagonal pores (20.3 ± 3.2 kPa), rhombic pores with 34° (28.7 ± 4.9 kPa) and 70° (73.0 ± 21.5 kPa) winding angles, outperforming gelMA-only controls (5.2 ± 2.1 kPa). Overall, the strongest reinforcement in terms of Young’s Modulus was provided by the 70° rhombic structure (10.8 ± 3.3 kPa) when compared to 34° rhombic (3.5 ± 0.3 kPa) and hexagonal (3.2 ± 1.2 kPa) structures (Figure 3E). The difference is likely caused by the specific pore sizes present in these MEW constructs as well as the compliance of the different repeating compartments in deforming along with the hydrogel during the tensile displacement. The effect of pore size of MEW meshes has already been characterized in previous studies.^[25, 43]^ The hexagonal constructs feature a higher stiffness compared to the rhomboid structures in radial deformation as an effect of their geometry, which is likely the main cause for the linear rise of stress in the recorded measurements and is supported by other studies on the influence of MEW geometries on their mechanical behavior.^[44]^ The rhomboid structures illustrated a certain degree of flexibility depending on the winding angle, allowing the construct to behave elastically in the direction of radial tensile deformation before transitioning to plastic deformation. This allows the rhomboid geometry to assume vastly different mechanical properties depending on the chosen winding angle.^[45]^ The aforementioned effect is presented in the recorded difference between the 34° (Peak Stress 28.7 ± 4.9 kPa / Young’s Modulus 3.5 ± 0.3 kPa) and 70° (Peak Stress 73.0 ± 21.5 kPa / Young’s Modulus 10.8 ± 3.3 kPa) rhomboid orientations (Figure 3D). The deviation between the generated samples is minimal within their group (apart from the 70° rhomboid constructs), highlighting the stable manufacturing process of the VolMEW constructs. The hexagonal structures differ from the rhomboids in their elastic properties due to the higher number of crossover points and the overall stable hexagonal geometry, resulting in a nearly linear mechanical behavior over the entire displacement range. For biological applications connective tissues and blood vessel walls are typically characterized by tensile stress-strain curves with an initial toe region (Supplementary Figure S5), indicative of the gradual recruitment of the ECM fibers in the direction of the application of the stress, followed by a stiffer region at higher deformations. A similar profile can be obtained with the rhombic MEW reinforcements at low winding angles (34°), while stiffer meshes in the radial direction can be obtained at higher winding angles (70°).^[45]^ To enable a comparison to natural tissues, two porcine coronary arteries were measured in the same fashion as the VolMEW constructs. When comparing the rhomboid constructs to the porcine coronary arteries, the 70° (Blood Vessel 1: 72.4 kPa / 70°: 75.5 kPa) and 34° (Blood Vessel 2: 38.7 kPa / 34°: 37.3 kPa) VolMEW constructs showed a good approximation of the maximum stress levels, while the 34° rhomboid reinforced constructs also showed a comparable overall curve trajectory to the physiological specimens (Figure 3F). Due to inter-patient variability and the wide range of mechanical properties that vessels display even when taken from adjacent anatomical locations,^[46]^ these results underscore the versatility of the proposed VolMEW system to modulate the mechanical profile of the printed composite tubes in order to approximate physiological blood vessel mechanics and to utilize the mesh design to account for natural variation.

Based on these findings, combined with the previously established superior light attenuation performance, further experimentation focused on the 34° rhombus geometry. A major benefit of MEW is the highly organized manner of fabrication and the consistent stacking of fibers, enabling a high degree of reproducibility and an additional adjustment point for mechanical properties.^[47, 48]^ The evaluation of the effect of different layer heights for the 34° rhomboid constructs illustrated an increase of maximum recorded stress (20L: 18.5 ± 4.2 kPa / 40L: 34.3 ± 3.3 kPa / 60L: 37.3 ± 1.5 kPa), as well as an increase of Young’s Modulus. (20L: 2.9 ± 0.3 kPa / 40L: 3.3 ± 1.1 kPa / 60L: 5.3 ± 0.9 kPa) (Figure 3G, H). Assessing the ability of the constructs to operate in a range of strains closer to biological values over an extended period is a valuable metric to record when looking into the mechanical properties of tubular constructs. A 200-cycle displacement to 20 % strain was used, and it revealed an increase in stiffness as a function of the number of layers of the MEW reinforcement geometries (control: 1 ± 0.3 kPa / 20L: 3.6 ± 0.2 kPa / 40L: 3.8 ± 1 kPa / 60L: 6.0 ± 1.3 kPa), except for the difference between 20 and 40 layers, where the SD and limit of recording resolution prevented any discernible significance from being recorded (Supplementary Figure S5C). It should be noted that although we focused on PCL as the MEW platform material in this study, the mechanical properties of the MEW fiber reinforcements could also be tuned by utilizing different biomaterial inks.^[49–51]^

In addition to tensile testing and burst pressure analyses, the shape stability of these VolMEW cylinders was assessed by evaluating their bending resistance. While a degree of bending flexibility can be desirable to manipulate the VolMEW tubes, unwanted collapse due to the inability of the tubes to sustain their own weight can be detrimental. The experiment was set up in a manner similar to the previously proposed filament collapse test, which has been used to evaluate the filament stability of extrusion-based printing materials.^[52]^ Briefly, long (27.5 mm) composite tubes (t_print_ = 16.4 s) were placed to bridge the distance across two columns spaced at increasing distance from each other (1 – 25 mm), and the gravity-induced flexural deformation of the tube was imaged and measured, as a function of the layer height (34° rhombic geometry) (Figure 3I, J). The most striking effect of the reinforcing MEW scaffolds was observed at the longest spanning distance of 25 mm. At this gap distance, where meshless 8% (w/v) gelMA cylinders completely collapse over the entire height of the column structure, increased resistance to bending is observed in the hybrid VolMEW samples as layer height increases (Figure 3I). Spanning the largest gap of 25 mm, the 60-layer construct only underwent an average maximum deformation distance of 1.16 ± 0.04 mm, significantly less than the 40- and 20-layer constructs (1.85 ± 0.05 mm and 4.09 ± 0.38 mm respectively). A similar trend in bending resistance was also observed for some of the smaller spanning distances (20, 16 and 12 mm), down to the 8 mm gap and below, where the maximum deformation becomes undetectable across all samples (Figure 3J). Overall, these mechanical evaluations confirm that the convergence of MEW with VBP results in hybrid constructs exhibiting superior mechanical stability compared to pure gelMA constructs, thus enabling applications of these hydrogel-based constructs in a broader array of biologically relevant settings.

With a thorough understanding of the mechanical reinforcement provided by MEW fibers in the established VolMEW converged approach, the unique advantages of the VP process were further explored to create advanced and geometrically complex tubular structures (Figure 4). Attempting to introduce cell-laden hydrogels within tubular porous structures, such as those produced by fused deposition modelling, melt electrowriting or solution electrospinning could be simply performed via casting in tubular molds. A major drawback of such casting approaches is the lack of precise control, which only allows the production of homogeneous cylindrical tubes consisting of a single gel layer (and thus a single layer of encapsulated cells). Alternative approaches are therefore needed to obtain, for instance, multilayered walls analogous to those found in vivo in vessels larger than 100 µm in diameter. With the positional accuracy provided by VP, the relative orientation of the hydrogel layers in relation to the MEW mesh can be freely designed and modified (Figure 4A). This was demonstrated with prints in which the hydrogel layer reached the center of the mesh, leaving either the outer region of the scaffold gel-free (inner print; Figure 4Ai, iii, v) or its inner region gel-free (outer print; Figure 4Aii, iv, vi). Such structures would have potential applications in vascular tissue engineering by allowing a second layer of gel to be printed over the exposed region of the scaffold to guide cell attachment and directionality,^[28, 53]^ or to incorporate multiple materials and cell types within the same construct. Another physiologically relevant application of the positional control of hydrogel printing within the MEW mesh is the creation of micron-scale fenestrations (Figure 4B), such as those commonly found in permeable vessels throughout the human body. These fenestrated vessel walls, already present in neovascular structures during development,^[54]^ are abundant in different regions of the body where permeability is critical for nutrient and waste exchange, including the renal glomerulus,^[55]^ intestinal blood vessels,^[56]^ the nasal mucosa,^[57]^ and are present in different areas of the central nervous system,^[58]^ including the blood-brain barrier in certain neurological diseases including strokes.^[59–61]^ Apart from these naturally occurring fenestrated structures, the clinical application of fenestrated stents for the repair of damaged vessels and aneurysms has also been a topic of interest, given the need to maintain transmembrane transport in the repaired vascular structures.^[62–64]^ In this study, we have demonstrated how to generate minute fenestrations by taking advantage of the natural interactions at the interface between the bioprinted hydrogel structures and the MEW mesh (Figure 4C). During the fabrication of MEW scaffolds, intersecting print paths of the same layer (crossover points) create a local elevation due to more material being stacked on top of each other, while at the same time there is a sagging effect between two crossover points that further amplifies the elevation effect. These crossover points are herein exploited to create fenestrated structures (Figure 4Ci). To achieve this, hollow gelMA cylinders were volumetrically printed exactly in the center of a MEW scaffold, leaving the inner base of the mesh and the tip of the crossover points exposed, allowing for small fenestrations (≈10-micron range, comparable to the MEW fiber thickness) to form on the outer surface of the hydrogel layer (Figure 4Cii, iii). These small fenestrations were successfully produced throughout long tubular structures and remained perfusable, as evident by the outflow of Alcian Blue stain through the exposed crossover points (Figure 4D). Importantly, the number and distribution of these fenestrations can be controlled by adjusting the number of pivot points of the MEW tubes, an easily adjustable parameter in the code used to fabricate these tubular scaffolds. It is clearly visible that the 4-pivot point scaffold (Figure 4Di) yields fewer fenestration regions compared to the 8-pivot point scaffolds (Figure 4Dii). The VP design alone was also tested to fabricate fenestration-like structures of decreasing size by stacking cylinders on top of each other with a minimum reproducible gap size of 464 ± 48 µm (Figure 4Ei). These cylinders remained bound together by the interwoven MEW fibers and also allowed lateral Alcian Blue outflow (Figure 4Eii). The combination of macroscale VP- and micron-scale crossover VolMEW fenestrations (as low as hundreds of nanometers)^[48]^ could introduce hierarchical pore distribution with spatial control along the length of the tube, while spanning a wide range of pore sizes within the same hydrogel shell. It should be noted that the density of larger, VP-induced pores should be carefully controlled, as placing too many of these gaps in close proximity to each other could significantly affect the structural stability of the printed constructs. These experiments further demonstrate the potential of the VolMEW convergence, as the incorporation of MEW constructs not only plays a mechanical role in VP printing, but can also be exploited to create more complex, native-like tubular structures when combined with the design freedom offered by volumetric printing.

**Figure 4:**
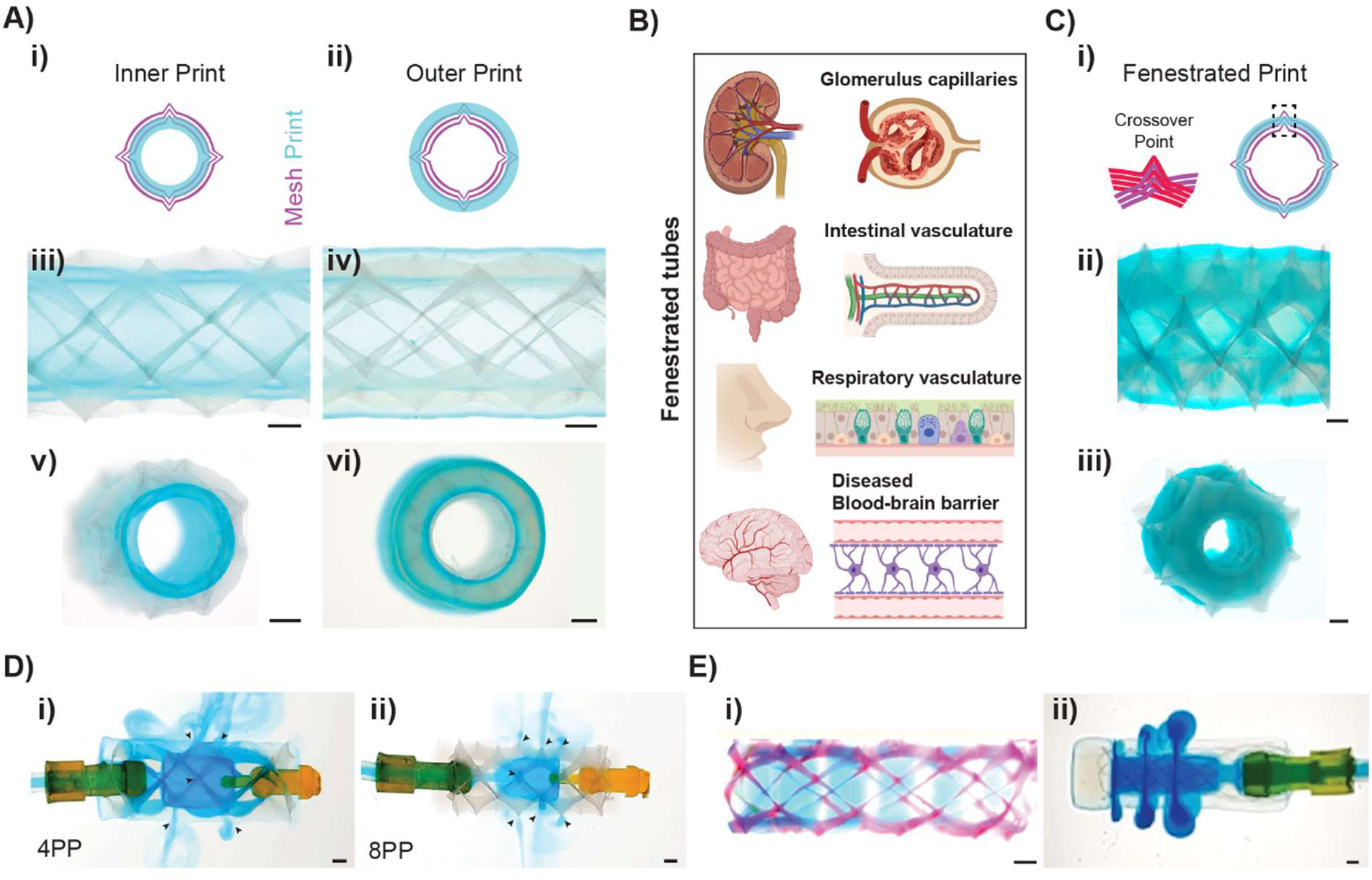
Printing of complex tubular structures and fenestrations. A) Graphical overview of possible i) inner and ii) outer printing strategies, in which the hydrogel embeds only the inner or outer region of the MEW construct respectively, leaving the rest exposed. Stereomicroscopic images of these iii, v) inner and iv, vi) outer prints from iii, iv) top and v, vi) side views. B) Various native tissues/structures have fenestrated structures to enhance transport or signal transduction, such as blood vessels in the renal glomerulus, intestinal vasculature, the nasal mucosa, and various regions of the central nervous system such as the blood-brain-barrier (Created with BioRender.com). C) VolMEW printing of fenestrated tubular constructs i) Graphical representation of the characteristic crossover points of tubular MEW meshes, and the resulting fenestrations when a thin layer of bioresin is printed exactly in the middle of the MEW construct. Stereomicroscopic images of the ii) top and iii) side views of such fenestrated structures. D) MEW induced fenestration showing local filtration at crossover points where the mesh pierces the hydrogel construct with i, ii) adjustable leakage over time based on the number of crossover points (black arrows) controlled by pivot points (PP). E) Circular fenestrations i) lightsheet maximum projection highlighting the ability to tailor fenestration size ii) light microscope image of the perfusable VolMEW construct with controlled leakage of Alcian Blue stain through the fenestrations, undisturbed by the presence of the MEW meshes (scale bars = 1 mm).

In addition to introducing the freedom to control the degree of alignment and the presence of pores and fenestrations, the hybrid VolMEW approach also facilitates, for the first time, the integration of complex architectural structures in the outer and inner regions of the MEW-reinforced tubular structures (Figure 5). Current approaches, based on hydrogel casting and extrusion bioprinting do not easily allow for the integration of complex architectural elements around and within the pre-existing MEW mesh, mainly due to the presence of moving needles in extrusion printing, and the challenges of mold removal in the case of casting. By fully encapsulating the MEW scaffold in a single step using the VP approach, hydrogel regions with increased design freedom were created. To demonstrate the addition of complex geometries on the outer side of the VolMEW tubes, branching channels stemming from the reinforced tube were successfully printed (t_print_ = 18 s). Both the reinforced and non-reinforced channels were homogeneously perfused, as the porous MEW mesh allowed for a seamless connection between the channels, demonstrating the potential to create more complex and reinforced branching structures that are crucial to replicating native vascular networks (Figure 5A). Additionally, the ability to print complex structures within the lumen of VolMEW reinforced tubes was demonstrated, a mathematically derived Schwarz D lattice was printed within the gelMA cylinder, as proof-of-concept design (Figure 5B). While these types of structures have previously been printed using a volumetric printer,^[33]^ it is remarkable that, even in the presence of the opaque MEW mesh, these fully perfusable structures could be resolved with a printing accuracy of 764 ± 48 µm (t_print_ = 14.8 s). Further applied examples of even more complex structures, functional components recurrent in blood vessel types, were successfully printed inside the VolMEW tubes. A simplified venous valve model (Figure 5Ci) was successfully printed at high resolution within VolMEW reinforced tubes, with leaflets measuring 232 ± 10 µm (t_print_ = 11.9 s) (Figure 5Cii). The valve, as found in the native structure, was able to control flow in a unidirectional manner in response to flow pressure, with the leaflets bending towards each other and closing, preventing flow in one direction (Figure 5Ciii), but remaining unstressed and open in the other direction, allowing uninterrupted flow (Figure 5Civ).

**Figure 5:**
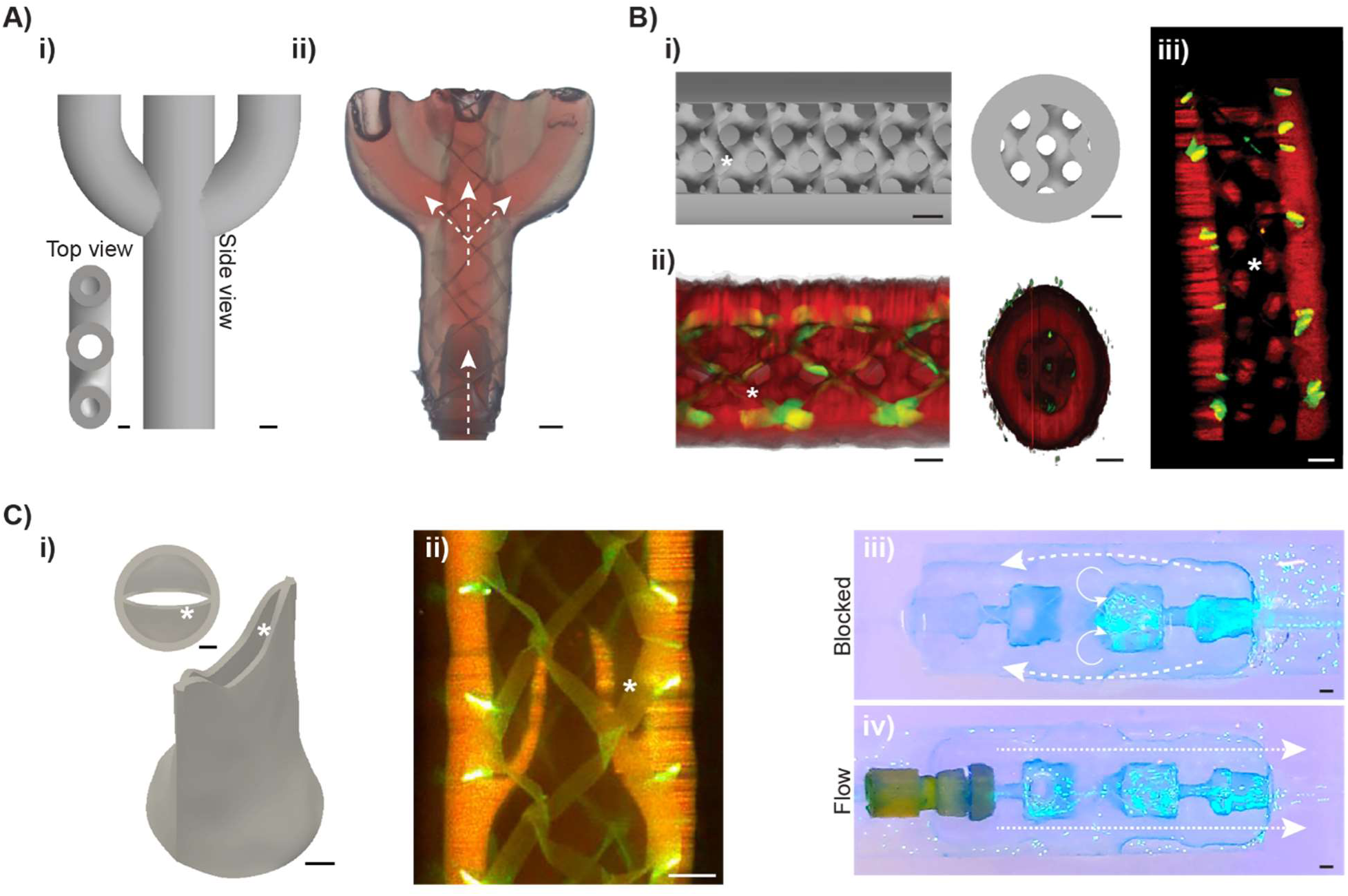
High-resolution VolMEW printing demonstrating perfusable external and internal features of physiological relevance. Ai) Digital model of a branched structure and ii) stereomicroscope images of the printed structure being perfused in a homogeneous manner (arrows indicate the direction of flow). Bi) A digital model of a VolMEW printed Schwarz D structure inside a cylinder containing the reinforcing MEW mesh with ii) its corresponding lightsheet 3D volume reconstruction and iii) a central lightsheet fluorescence image of the printed structure with the gel structure (red) and the encapsulated MEW mesh (green) highlighting the high printing resolution within the inner diameter of the mesh. Ci) Digital model of a simplified venous valve model and ii) a central lightsheet section of the printed construct showing the high resolution leaflets (*) of the valve. iii) Digital images of fluorescent beads perfused through the valve structure showing unidirectional flow (dotted lines show the outline of the printed construct and arrows indicate the direction of flow). (scale bars = 1mm).

After successfully demonstrating the ability to print these different geometric features (valves and complex branched structures) and given the promising mechanical properties of the hybrid VolMEW scaffolds, the possibility of forming cell-laden constructs containing multiple cell types by both direct bioprinting and by post-print seeding was investigated. To make this possible, we devised a strategy to sequentially, volumetrically print multiple cell-laden hydrogel layers (Figure 6A), followed by a seeding phase in the lumen of the tube, resulting in a three-layer construct (Figure 6B). As preparatory step, using a single fluorescently labeled suspension of human mesenchymal stromal cells (DiD-hMSCs), it was shown that cell-laden gelMA resins could be printed to precisely envelop the MEW scaffold (t_print_ = 14 s), and the resulting cellularized construct could then be seeded with human umbilical vein endothelial cells (HUVECs) after printing (Figure 6C). This resulted in a two-layer construct with independent cell regions, an inner HUVEC layer (Figure 6Di) and a 3D gelMA layer encapsulating hMSCs and the MEW scaffold to provide mechanical stability to the structure (Figure 6Dii). Next, as a proof-of-concept, a “protovessel” consisting of three distinct cellular layers was fabricated using the hybrid VolMEW approach. Native macro-scale vessels, such as veins or arteries, have three distinct cell layers, the tunica intima, media, and externa, each consisting of different cell types and matrix composition required for proper vessel function (Figure 6E). This tubular construct consisted of the previously described two-layer approach, but by adding a second VBP step with a new cell suspension (hMSCs labeled with a different fluorescent dye, Dil), an outer cell and material layer was successfully printed, creating a three-layer architecture that approximates the layered structure of a native vessel (Figure 6F, G). Furthermore, the possibility of creating fenestrated, cellularized protovessels was also demonstrated using the principles described in Figure 4. By first creating the first hydrogel layer as an inner print, leaving the outer edge of the vessel exposed, the second layer could still be printed within the MEW scaffold, leaving the small crossover points of the mesh exposed to the outside, creating the previously shown, perfusable fenestrations in a cellularized model (Figure 6H). These final proof-of-principle prints demonstrate that the use of the VolMEW hybrid approach can be exploited in several ways to increase the complexity of current tissue engineered macrovascular-inspired structures. Several biofabrication approaches have been explored to create architecturally complex vessels, including sacrificial extrusion templating,^[65, 66]^ coaxial extrusion bioprinting,^[67]^ suspended bioprinting^[68]^ and suspended sacrificial printing,^[69]^ digital light projection printing,^[70–72]^ and acoustic wave patterning.^[73, 74]^ Expanding on this toolkit, VolMEW introduces the ability to both tune the mechanical properties of the construct and to introduce custom-designed patterns of microfibers, which have been used in previous work to facilitate stromal cell alignment.^[31]^ Notably, and with broad applications beyond of vascular-mimetic printing, VolMEW allows to freely sculpt the hydrogel component, creating features that intertwine with the MEW mesh, or, for example, incorporate fenestrations and branches relevant for hierarchical networks,^[75]^ and valve-like structures, of relevance to regulate flow and local pressure in vessels and more generally in fluidic components. These architectures can be printed in seconds with no need for support materials, incorporating a mechanically reinforcing MEW mesh without compromising print resolution. Overall, these features make VolMEW promising for the creation of next-generation tubular grafts with customizable designs that can be tailored to tissue-specific requirements and produced in high-throughput.

**Figure 6:**
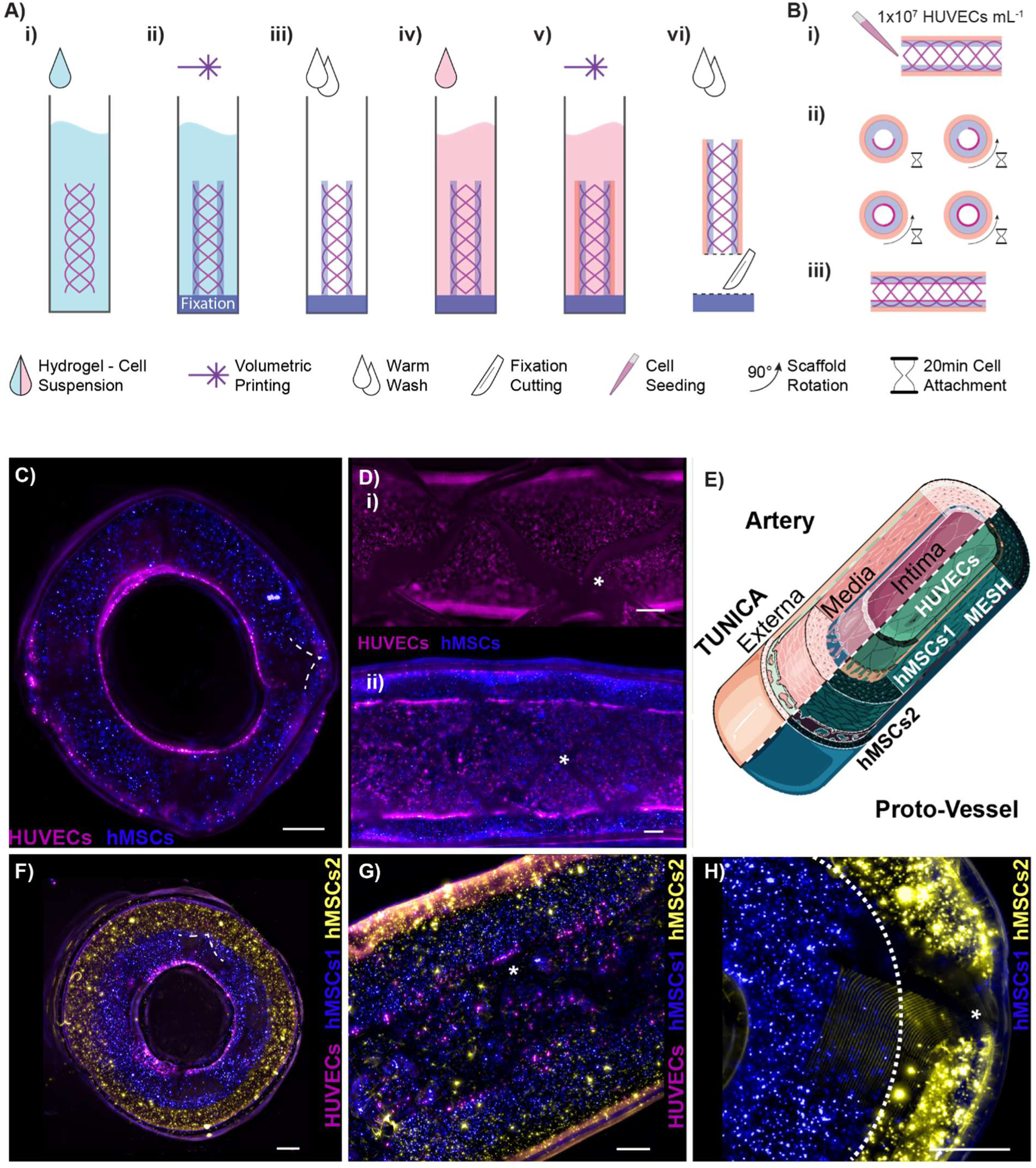
Sequential VolMEW printing of cell-laden, multi-material and multi-layer tubular constructs. A) Graphical overview of the multi-material VolMEW printing process with cell-laden bioresins. i) The MEW mesh is inserted as previously described followed by ii) VP printing of the scaffold with an overexposed fixation, whereby the construct is firmly attached to the printing vial. After iii) washing the residual, uncrosslinked hydrogel, iv, v) the process is repeated with the second material. Finally, the vi) fixation is cut with a scalpel. B) Depiction of the HUVEC seeding process, where first i) 1x10^7^ cells mL^-1^ are pipetted into the construct, followed by ii) 20 minutes of incubation periods ending with 90° rotations. The process is repeated four times to homogeneously cover all sides to create iii) the final three-layer VolMEW construct. C) Perpendicular and Dii) longitudinal cross-sectional fluorescence images of a two-layer VolMEW-printed tubular construct consisting of VBP-printed hMSCs (blue) in the gel layer encapsulating the MEW mesh and i) a HUVEC-seeded lumen (magenta). E) Diagram of native vessel structures compared to the VolMEW printed proto-vessels. F) Perpendicular and G) longitudinal cross-sectional fluorescence images of a tree-layer VolMEW printed tubular construct consisting of VBP-printed hMSCs (blue and yellow) in the gel layer encapsulating the MEW mesh and a HUVEC-seeded lumen (magenta). H) Sequentially printed multi-material construct with crossover points exposed, embedding the MEW mesh in both layers The Figure E) was partly generated using Servier Medical Art, provided by Servier, licensed under a Creative Commons Attribution 3.0 unported license. (C-H: mesh indicated; scale bars = 500 µm).

## CONCLUSIONS

This study demonstrates, for the first time, the convergence of volumetric bioprinting with melt electrowriting to build geometrically complex objects with enhanced mechanical properties, even when using low stiffness bioresins commonly used in VP. In the first stage of the study, we presented the VolMEW setup, which takes advantage of the thermal gelation properties of gelMA and a custom-designed guide system to precisely place and align MEW constructs of different architectures. While the presence of MEW constructs in the VP vial is shown to attenuate the tomographic light path, VP-printed layers were successfully, and precisely sculpted onto and across the opaque microfibrous meshes with high shape fidelity. Furthermore, the reinforcing effects observed in these converged printing constructs show mechanical advantages in flexural, burst and tensile strength compared to non-reinforced scaffolds, reducing the gap between these hybrid biofabricated constructs and native tubular tissues of biological relevance, such as vascular structures. This newly developed approach also retains the high printing speeds (< 20 seconds) and unparalleled design freedom associated with conventional VP. Exploiting this architectural freedom, an array of hierarchical, physiologically relevant constructs could be produced. This includes custom-designed and distributed wall fenestrations and complex printing of gyroid-structures, bifurcated channels and a functional venous valve model within the reinforced hydrogel tubes. As a proof-of-concept, the possibility of creating multi-material and multi-cellular structures through a sequential printing approach was also demonstrated. The three distinct layers found in native macro vessels (i.e., veins and arteries) could be replicated in these reinforced, composite structures, demonstrating the potential to create hierarchical living constructs with the VolMEW approach. Notably, since the MEW meshes can be bulk produced and stored to printing, future applications can be envisioned, in which off-the-shelf mesh geometries could be readily loaded into a volumetric printer, to add the hydrogel and cellular components, just before their intended application. Overall, this novel technique poses the possibility of creating volumetric composite objects from materials with very different chemical and physical properties (hydrogels and thermoplastics) with enhanced mechanical properties and high design freedom. By leveraging the advantages of the MEW and VP technologies in the present approach, these findings also provide exciting opportunities for future hybrid applications with other opaque materials (i.e., ceramics, metals) for advanced tissue engineering strategies.

## EXPERIMENTAL SECTION

Materials: For melt electrowriting, medical-grade polycaprolactone (PCL) (PURASORB PC 12, Corbion Inc., Gorinchem, The Netherlands) was used to fabricate the MEW tubular scaffolds. As a bioresin for volumetric (bio)printing, gelatin methacryloyl (93.5% DoF) was synthesized as previously reported^[76]^ and used as 5% w/v (cellular experiments), 15% w/v (venous valve) and 8% w/v (all others) solution in phosphate-buffered saline (PBS). As photoinitiator, 0.1% w/v lithium phenyl(2,4,6-trimethylbenzoyl)phosphinate (LAP, Tokyo Chemical Industry, Japan) was supplemented to the hydrogel precursor solution to initiate the photocrosslinking reaction.

Melt Electrowriting of Tubular Structures with Different Geometries: Tubular MEW constructs were processed using two custom-made melt electrowriting devices with a cylindrical and interchangeable collector. One device, used to fabricate tensile test specimens, employs an Aerotech axis system (PRO115) and the A3200 (Aerotech, USA) software suite as coding and machine operating interface. Polypropylene cartridges, and 22G flat-tip needles (Nordson EFD, USA) were used. The second device, used for all other experiments, has been described elsewhere^[28]^ and was used with an electrically heated 3 mL glass syringe and a 25G needle. A modified code was developed similar to previous work,^[25]^ to move the collector in translational and rotational directions for precise fiber placement onto a rotating steel mandrel at predetermined winding angles.

Thermally Controlled Incorporation of Tubular MEW Meshes into Volumetric Printing Setup: In order to insert the tubular MEW scaffolds in a perfectly centered and reproducible manner, the scaffolds were transferred to 3mm pultruded carbon fiber rods (easycomposites, The Netherlands) matching the inner diameter of the MEW mesh. To center the rod inside a Ø10 mm cylindrical borosilicate glass vial compatible for use with the volumetric printer, small inserts were printed on a Perfactory 3 digital light projection (DLP) 3D printer (Envisiontec, Germany) using PIC100 resin to fit the rod and perfectly match the inner diameter of the vial. To fix the samples in the center of the vial, the tubular mesh was placed on the end of the insertion rod with 5mm offset. The printing vial was then filled with preheated (37°C) bioresin and the rod was inserted. The vial was subsequently placed in ice water up to the beginning of the insertion rod until gelation occurred. The gelled bioresin holds the tubular mesh firmly in place. The insertion rod is raised to the upper limit of the tubular mesh and the gelation process repeated, resulting in a centered sample.

Volumetric (Bio)Printing Process: A commercial Tomolite v1.0 (Readily3D SA, Switzerland) volumetric printing setup was used to fabricate the hybrid VolMEW constructs. The gelMA bioresin-infused MEW meshes were prepared as described above in Ø10 mm cylindrical borosilicate glass vials and kept cool at 4° C to ensure that the resin remained thermally gelled. Custom-designed CAD files (Fusion360) were loaded and processed using the Readily3D Apparite software (b11409a). The average light intensity across all prints was set at 12.16 mW/cm². Venous valve models were additionally printed using Apparite’s proprietary settings for a bulk length of 400 µm and a bulk multiplier of 2x. After printing, the thermally gelled bioresin was gently washed with pre-warmed PBS at 37° C to remove the uncrosslinked resin from the printed structure. To achieve homogeneous crosslinking and facilitate sample handling, the samples were immersed in a 0.01% w/v LAP solution and post-cured for 5 min in an UV oven (Cl-1000, Ultraviolet Crosslinker, λ = 365 nm, I = 8mW/ cm^2^ UVP, USA).

Measurements of MEW Mesh Light Attenuation Effect during VP: Tubular MEW meshes were placed in cylindrical vials containing the gelMA resin. The vials were then placed within the print area of the volumetric printer and coupled to the rotary stage (Newmark RB-90). Index matching was achieved by immersing the vial in a square cuvette containing distilled water. A virtual beam was generated by encoding a circular region of on-sate pixels on a DMD (Vialux Hi-Speed V-7000) illuminated by a collimated 520nm source. This corresponded to a beam waist diameter of 200 ± 10 µm. The vial was positioned so that the virtual beam was incident on the central axis of the embedded MEW mesh and the vial. A biased photodetector (Thorlabs DET36A2) was positioned 100 ± 5 mm behind the vial and was used to measure the relative intensity of the incident light after attenuation by the scaffold. A digital acquisition board (National Instruments USB-6001) was used to acquire the voltage signal from the photodiode. In order to perform the measurement, the virtual beam was turned on and the vial was rotated at a rate of 36 deg sec^-1^ over a full rotation. The resulting voltage signal from the photodiode was recorded at a sampling rate of 50 Hz. For each sample, this measurement was acquired at 3 locations on the MEW mesh, with the virtual beam being offset by +3 mm and -3 mm from its central position. The acquired data was normalized with 100% being the maximum amount of light detected across all runs. The runs were then rotated to start at the first peak using Savitzky-Golay filtering, followed by finding the first local maximum, and the resulting phase-matched raw data were averaged using time synchronous averaging (Matlab R2022a). The area under the curve was approximated using discrete trapezoidal numerical integration and the data were normalised (deg) and plotted (Matlab R2022a).

Tensile Testing of Hybrid VolMEW Constructs: To determine the radial mechanical properties of the VolMEW constructs, a customized two-pin mounting setup was used on a dynamic mechanical tester (Electron-Force 5500, TA Instruments, USA). Two metal pins were inserted through the luminal cavity of the constructs and a radial tensile force was applied during the test procedure (Figure 3A). Samples were measured in a 100-cycle waveform setup with a peak displacement of 18% strain with respect to the inner tube diameter. Construct measurements were evaluated after the initial hysteresis had subsided and the peak force had stabilized over several cycles. A second evaluation was a pull to 166% strain to elucidate the maximum stress values at beyond physiological values of small diameter blood vessels.

Burst Pressure Analysis of VolMEW Constructs: A Vieweg DC 200 (Vieweg, Germany) dispenser was used with a custom tubing array that included a pressure gauge and a Luer connector to a custom-made bioreactor assembly part created by DLP printing on a Prusa SL1s resin printing system (Prusa, Czech Republic) with Dreve FotoDent guide 405 nm (Dreve, Germany). The resin was also used to fix VolMEW constructs liquid tight into the assembly part, a thin cover of FotoDent has been applied to the connectors and cured with a Prusa CW1 curing and washing station (Prusa, Czech Republic). Vaseline was then injected into the VolMEW construct through an attached printer cartridge (Nordson, USA) until failure of the constructs could be observed and burst pressure was recorded. Data collection was done by digital video recording of VolMEW constructs consisting of MEW meshes of different architectures and number of layers.

Bending Resistance Evaluation of VolMEW constructs: To evaluate the effect of MEW mesh incorporation on the bending resistance of the hydrogel constructs, 27.5 mm long VolMEW scaffolds were printed with meshes of rhombic pores with 34° winding angle consisting of different layer numbers (0, 20, 40 and 60). Based on the previously established filament collapse test developed to evaluate the shape fidelity of bioinks^[52]^ similar 5mm column structures were printed from polylactic acid (PLA, MakerPoint) using a fused deposition modeling printer (Ultimaker S3, Ultimaker, The Netherlands) with gaps of 25, 20, 16, 12, 8 and 4 mm, to assess the maximum deformation of the printed structures as they spanned each gap distance. Maximum deformation was measured as the lowest point the sample bent downwards from the top of the column structure.

Cell Isolation and Culture: Green fluorescent human umbilical vein endothelial cells (GFP-HUVECs; cAP-001GFP, Angioproteomie) were expanded in type I collagen precoated culture flasks in endothelial cell growth medium-2 (EGM-2) BulletKit medium (CC-3162, Lonza). Culture flasks were precoated with 50 μg ml^−1^ collagen I rat tail (354236, Corning) in 0.01 M HCl for 1 hour at 37°C, followed by two washes with phosphate-buffered saline (PBS). GFP-HUVECS were used in experiments at passage 6-7 and cultured in EGM-2 medium for expansion and differentiation. Human bone marrow-derived mesenchymal stromal cells (hbMSCs) were isolated from bone marrow aspirates of consenting patients, as previously described.^[77]^ Briefly, human bone marrow aspirates were obtained from the iliac crest of patients that were receiving spondylodesis or hip replacement surgery. Isolation and distribution were performed in accordance with protocols approved by the Biobank Research Ethics Committee (isolation 08-001, distribution protocol 18-739, University Medical Center Utrecht). Protocols used are in line with the principles embodied in the Declaration of Helsinki. MSCs were expanded in DMEM + GlutaMAX supplemented with FBS (10% v/v) and p/s 1%. The procedures for human tissue and cell isolation were approved by the Research Ethics Committee of the University Medical Center Utrecht.^[77]^ All cells were cultured at 37°C, 5% CO2 and used at passage 4-5. HbMSCs were cultured in DMEM + GlutaMAX supplemented with FBS (10% v/v) and p/s 1%. All cells were cultured at 37°C and 5% CO2 and used at passage 4-5.

Volumetric Bioprinting of Reinforced Multilayered Macrovessel-like Structures: Mechanically reinforced macrovessel structures mimicking the layered structure of native vessels (tunica intima, media and externa) were fabricated using a sequential bioprinting approach. The following procedure was used to fabricate two- and three-layer protovessel constructs. First, a 15mm hybrid VolMEW hollow tube was printed with 1 x 10^6^ cells mL^-1^ hbMSCs labeled with membrane staining DiD (ThermoFisher Scientific, The Netherlands) encapsulated in 5% w/v gelMA + 0.1% w/v LAP bioresin. In this first layer, a MEW scaffold (40 layers) with rhombic pores of 34° winding angle and 8 pivot points was incorporated and completely encapsulated by the printed gel layer of 1050 µm. For the 3-layer structure, the sample was washed after the first printing step and re-infused in a bioresin suspension containing 5 x 10^5^ cells mL^-1^ hbMSCs labeled with membrane staining DiI (ThermoFisher Scientific, The Netherlands), and a second outer layer of 800 µm was printed around the first cell layer. Samples were washed, post-cured for 5 minutes and cultured in hbMSC expansion medium overnight. The next day, the samples were placed on a rectangular mold with a concave slot for the tube to rest on, and the channels were seeded with a suspension of 1 x 10^7^ cells mL^-1^ GFP-HUVECs. Samples were rotated 90° every 20 minutes, for a total span of 80 minutes to achieve homogeneous cell seeding across the lumen of the protovessel. To create a fenestrated 3-layer protovessel, the same protocol was used, but the total width of the two cell layers was reduced to allow for overprinting of the MEW mesh outside the gel sample. Lateral and longitudinal cross-sections of the samples were imaged after 7 days of culture in EGM-2 BulletKit medium using a Thunder imaging system (Leica Microsystems, Germany).

Statistical Analyses: Results were reported as mean ± standard deviation (S.D.). Statistical analysis was performed using GraphPad Prism 9.0 (GraphPad Software, USA). For tensile test results, Origin 2022 (OriginLab, USA) was used. Comparisons between experimental groups were assessed via one or two-way ANOVAs, followed by post hoc Bonferroni correction to test differences between groups. Non-parametric tests were used when normality could not be assumed. Differences were considered significant when p < 0.05. Significance is expressed on graphs as follows: * p <= 0.05, ** p <= 0.01, *** p <= 0.001, **** p <= 0.0001.

## ACKNOWLEDGEMENTS

G.G., M.B.K., C.G. and P.N.B. contributed equally to this work. T.J. and R.L. contributed equally to this work. We would like to acknowledge Lisa Galaba, Sven Heilig and Philipp Stahlhut for their support in the experimental activities. This project received funding from the European Research Council (ERC) under the European Union’s Horizon 2020 research and innovation programme (grant agreement No. 949806, VOLUME-BIO). R.L. and J.M. acknowledge the funding from the ReumaNederland (LLP-12, LLP22, and 19-1-207 MINIJOINT). M.d.R., R.L. and J.M. acknowledge the funding from the Gravitation Program “Materials Driven Regeneration”, funded by the Netherlands Organization for Scientific Research (024.003.013). R.L. also acknowledges funding from the NWA-Ideeëngenerator programme of the Netherlands Organization for Scientific Research (NWA.1228.192.105). M.B.-K. and T.J. would like to thank the German Research Foundation (DFG, Project No. 326998133-TRR 225 – subproject Z01) for financial support. The DFG also supported the project with a “State Major Instrumentation Programme” (INST 105022/58-1 FUGG) that enabled SEM analysis of the samples. Further, M.B.K., C.G, J.G. and T.J. thank the European Union for support on printing strategies (European Fund for Regional Development – EFRE Bayern, Bio3D-Druck project 20-3400-2-10). T.J. acknowledges the European Union for funding via the European Union’s Horizon 2020 research and innovation program (BRAVE) under Grant Agreement No. 874827.

## Supporting Information

### Supplementary Figures

**Supplementary Figure S1:**
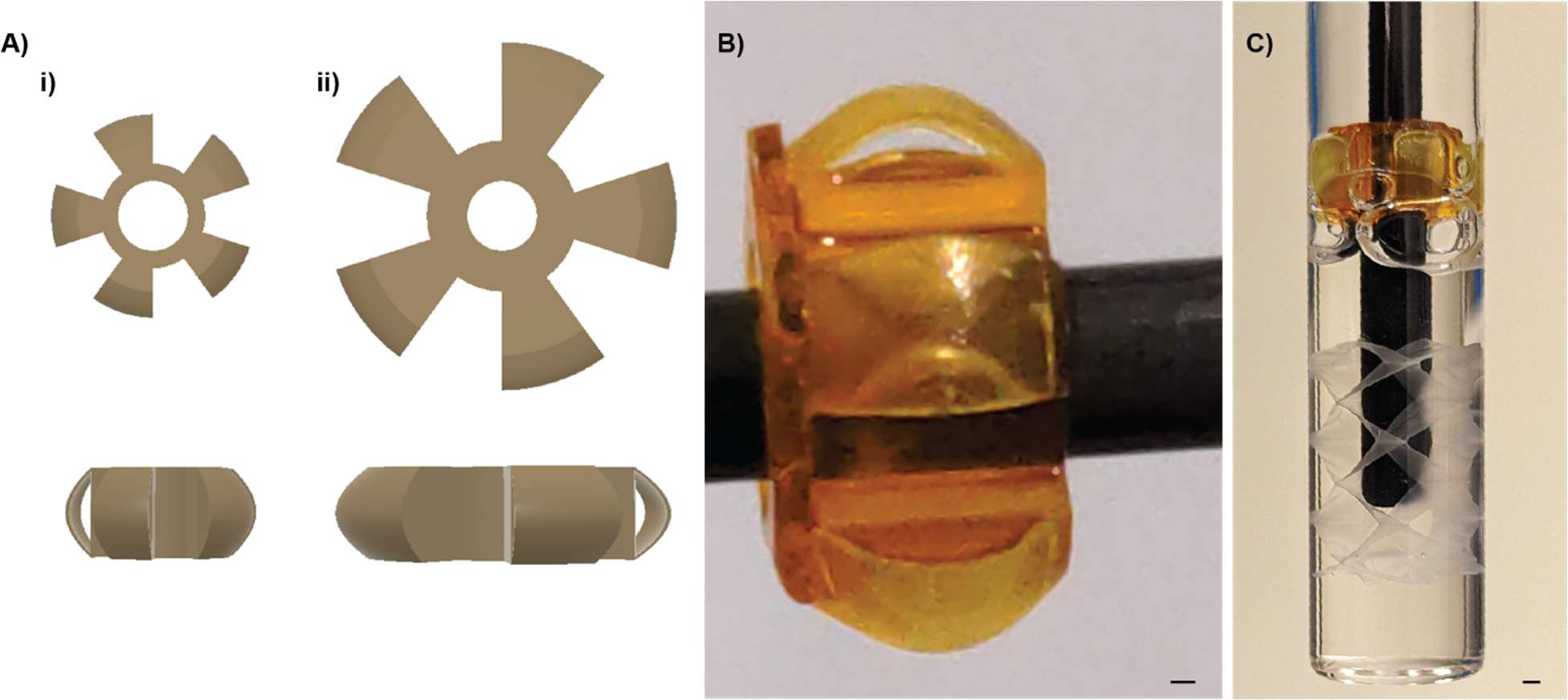
DLP printed alignment guides for tubular MEW meshes with 150 µm self centering wings. Ai,B) Winged guide for 10 mm and Aii) 20 mm Tomolite vials. B) Printed guide on carbon fiber rod for vial placement. C) Assembly with hydrogel in a vial prior to thermal gelation. (Scale bar = 1mm).

**Supplementary Figure S2:**
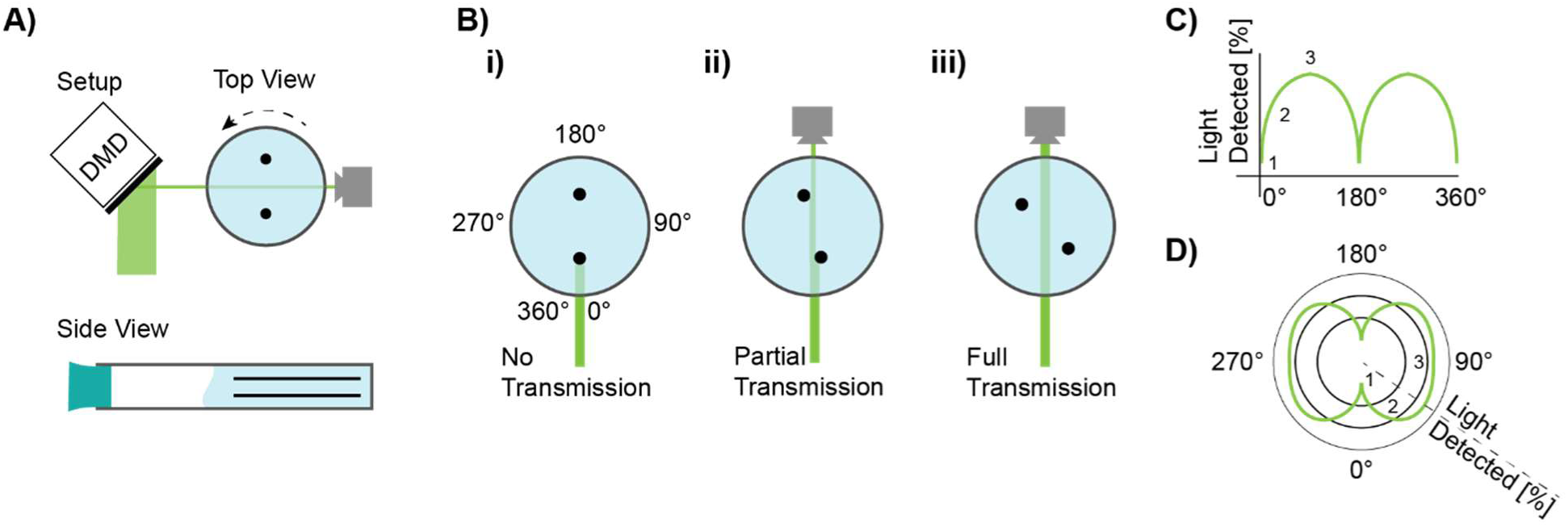
Overview of the light attenuation setup from measurement to polar plot. A) Setup with two hypothetical rods as attenuating objects. B) Transmission profiles of light with i) no transmission, ii) partial transmission and iii) full transmission. The corresponding C) Cartesian and D) polar plots.

**Supplementary Figure S3:**
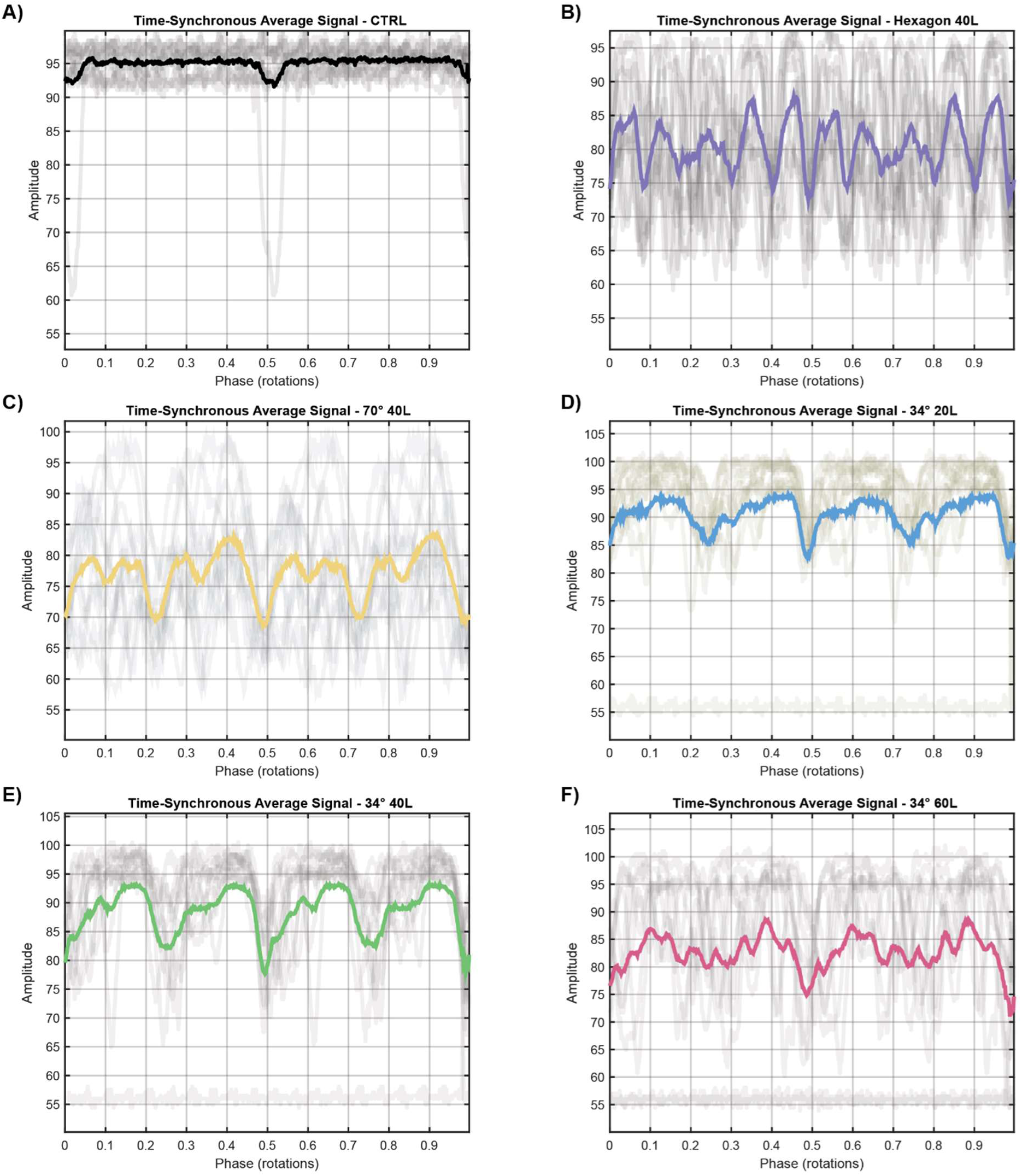
Time synchronous average (tsa) data with data set lines (gray) and highlighted tsa (color) for A) control, B) hexagonal (40L), C) 70° rhombus (40L), and 34° rhombi with D) 20-, E) 40-, and F) 60-layer heights.

**Supplementary Figure S4:**
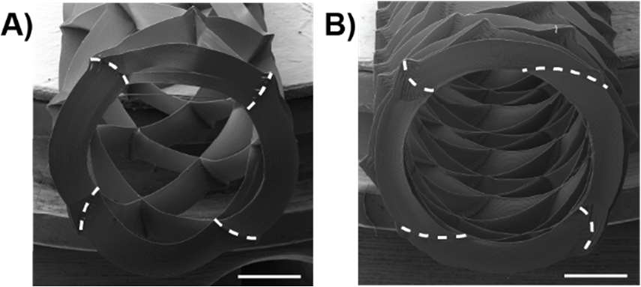
A) 34° and B) 70° winding-angle scaffolds with the uncompensated layer-shifting of crossover points with increasing wall height (dashed lines; scale bars = 1 mm).

**Supplementary Figure S5:**
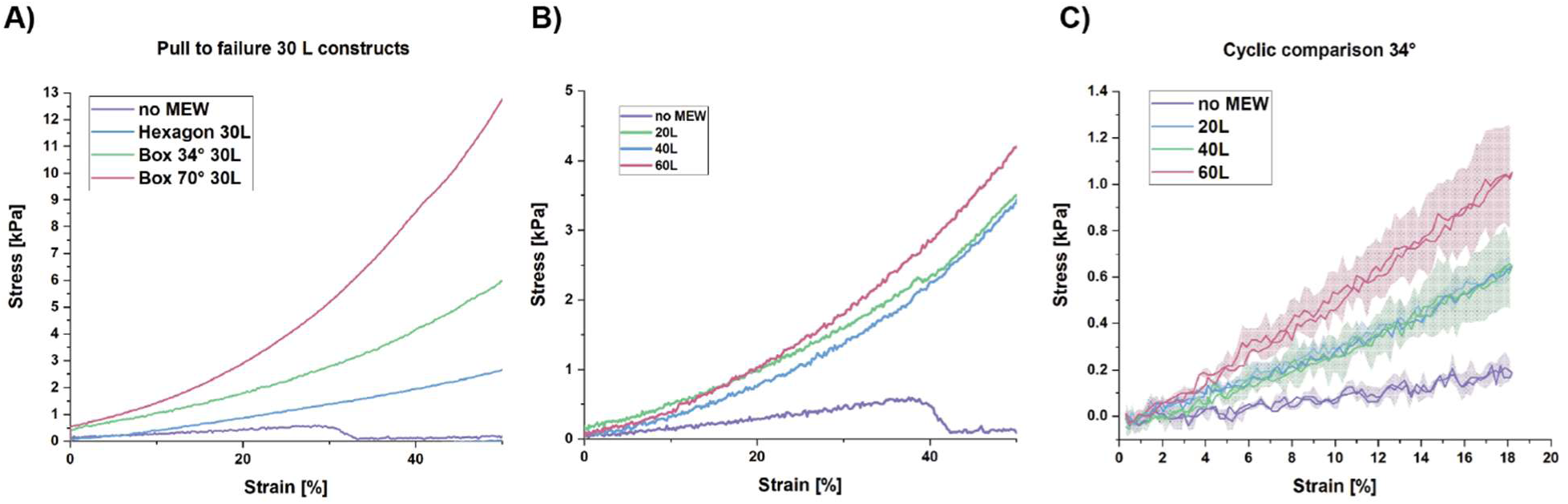
UTS zoom of the relevant Young’s Modulus region for A) different architectures and B) different layer heights. C) Cyclic radial stress-strain curves presented as their mean with SD values to illustrate the deviation within the same MEW reinforcement geometry.

